# Proteomic responses to ocean acidification in the brain of juvenile coral reef fish

**DOI:** 10.1101/2020.05.25.115527

**Authors:** Hin Hung Tsang, Megan Welch, Philip L. Munday, Timothy Ravasi, Celia Schunter

**Affiliations:** Swire Institute of Marine Science, School of Biological Sciences, The University of Hong Kong, Pokfulam, Hong Kong SAR; Australian Research Council Centre of Excellence for Coral Reef Studies, James Cook University, Townsville, Queensland, 4811, Australia; Marine Climate Change Unit, Okinawa Institute of Science and Technology Graduate University, 1919–1 Tancha, Onna-son, Okinawa 904–0495, Japan

**Keywords:** environmental proteomics, climate change, ocean acidification, behaviour, tolerance

## Abstract

Elevated CO_2_ levels predicted to occur by the end of the century can affect the physiology and behaviour of marine fishes. For one important survival mechanism, the response to chemical alarm cues from conspecifics, substantial among-individual variation in the extent of behavioural impairment when exposed to elevated CO_2_ has been observed in previous studies. Whole brain transcriptomic data has further emphasized the importance of parental phenotypic variation in the response of juvenile fish to elevated CO_2_. In this study, we investigate the genome-wide proteomic responses of this variation in the brain of 5-week old spiny damselfish, *Acanthochromis polyacanthus*. We compared the expression of proteins in the brains of juvenile *A. polyacanthus* from two different parental behavioural phenotypes (sensitive and tolerant) that had been experimentally exposed to short-term, long-term and inter-generational elevated CO_2_. Our results show differential expression of key proteins related to stress response and epigenetic markers with elevated CO_2_ exposure. Proteins related to neurological development were also differentially expressed particularly in the long-term developmental treatment, which might be critical for juvenile development. By contrast, exposure to elevated CO_2_ in the parental generation resulted in only three differentially expressed proteins in the offspring, revealing potential for inter-generational acclimation. Lastly, we found a distinct proteomic pattern in juveniles due to the behavioural sensitivity of parents to elevated CO_2_, even though the behaviour of the juvenile fish was impaired regardless of parental phenotype. Our data shows that developing juveniles are affected in their brain protein expression by elevated CO_2_, but the effect varies with the length of exposure as well as due to variation of parental phenotypes in the population.

## Introduction

Rising CO_2_ levels in the ocean are anticipated to affect the physiology and behaviour of marine organisms, with possible impacts on the structure and function of marine ecosystems (Wittmann and Pörtner, 2013; Cattano et al., 2018; Pörtner et al., 2019). Exposure to elevated CO_2_ has shown to alter important behaviours, such as navigation, habitat selection, learning and predator-prey interactions in some fish (Nagelkerken and Munday, 2016; Munday et al., 2019; Paula et al., 2019; Williams et al., 2019) and invertebrates (Watson et al., 2013; Jellison et al., 2016). For example, exposure to elevated CO_2_ impairs the natural avoidance behaviour of chemical alarm cues (CAC) in some fishes (Ferrari et al., 2011; Chivers et al., 2014; Welch et al., 2014; Ou et al., 2015; Welch and Munday, 2017; Laubenstein et al., 2019). Chemical alarm cues are compounds released from the skin of injured fishes, thereby providing a reliable cue of immediate predation threat (Ferrari et al., 2009). Most fishes respond to the presence of CAC from conspecifics by making behavioural decisions that reduce the risk of predation, such as reducing activity and seeking shelter (Ferrari et al., 2009). However, when exposed to elevate CO_2_, some fishes fail to exhibit behaviours that reduce their risk of predation. This impaired behaviour is hypothesized to be caused by the self-amplifying alteration in the chloride-bicarbonate current in the GABAA receptor in the brain, following acid-base regulation in reaction to elevated environmental CO_2_ (Nilsson et al., 2012; Chivers et al., 2014; Heuer et al., 2016; Schunter et al., 2019).

Studying the response to a change in environmental condition on the molecular level of an organisms can provide insight into the broad response of a tissue to this change. Some molecular processes might change to acclimate to the new condition and prevent the need of a change on the physiological, behavioral or whole organism level. On the other hand, some molecular patterns will be the underlying processes to the whole organisms’ physiological changes in response to the environmental alteration (Madeira et al., 2017). Transcriptomics can aid in understanding this molecular reaction whereas the protein response can elucidate possible post-translational modifications that have a more direct effect on cellular responses (López, 2007; Tang et al., 2015).

Previous brain transcriptomic studies on five-month-old spiny damselfish *A. polyacanthus* showed differential gene expression in GABA related pathways especially during short-term elevated CO_2_ exposure (Schunter et al., 2018). However, when elevated CO_2_ exposure was prolonged across generations, the transcriptional signal differs and shows an overexpression of genes related to sugar and energy production in the brain (Schunter et al., 2016), whereas the behavioural impairments persist (Welch et al., 2014). There is also large variation in the sensitivity of individual *A. polyacanthus* to moderately high CO_2_ levels (~700µatm), with a part of the population exhibiting some resistance to behavioural impairment (“tolerant” individuals) whereas others are behaviourally impaired when exposed to elevated CO_2_ (“sensitive” individuals) (Schunter et al., 2016). This variation in sensitivity to behavioural effects of elevated CO_2_ has a heritable component as fathers and their offspring show similar sensitivities after short-term exposure to elevated CO_2_ (Welch and Munday, 2017).

The protein signal of the tolerant and sensitive parental behavioural phenotype in the offspring has been previously investigated in five-month-old spiny damsel fish, by comparing control versus inter-generational CO_2_ exposed fish (Schunter et al., 2016). However, the early juvenile developmental period is one of the most critical life stages in fish (DePasquale et al., 2015) and environmental conditions could have strong impacts during this early life stage. Therefore, here we focus on investigating the proteomic responses of five-week-old juvenile *A. polyacanthus* exposed to elevated CO_2_ (750µatm) for different periods of time. In our experimental design we use the variation in the olfactory response behaviour of adult fish to study the influence of parental phenotype on brain protein expression of small juveniles. Further, these developing offspring of spiny damselfish from behaviourally tolerant and sensitive parents were exposed to different exposure lengths of elevated CO_2_: 1) Acute (5-days of elevated CO_2_), Developmental (5 weeks of elevated CO_2_ since hatching) and Inter-generational (previous parental exposure and 5 weeks of elevated CO_2_). We investigated the relevant molecular pathways involved in the reaction to the three different CO_2_ treatments by analyzing the differential expression of the brain proteome of 5-week old developing *A. polyacanthus*.

## Methods

### Experimental design

Adult *A. polyacanthus* were collected as described in Welch and Munday (2017) and exposed to elevated CO_2_ (754 ± 92 μatm), consistent with average atmospheric CO_2_ predicted by the end of the century according to the RCP6 emissions trajectory. After a seven-day elevated CO_2_ exposure, the reaction towards chemical alarm cues (CAC) in adults was tested as described in Welch & Munday (2017) to define the parental phenotype to be either tolerant (T; less than 30% of the trial time was spent in CAC) or sensitive (S; more than 70% of the trial time in CAC) (Figure 1). Individuals of the same behavioural phenotype (T or S) were paired into breeding pairs and either left at control condition (414 ± 46 μatm) or placed into elevated CO_2_ (754 ± 92 μatm) for three months until breeding started. Offspring clutches from each breeding pair were then placed into different experimental conditions resulting in a total of four treatment groups for each parental behavioural phenotype (T and S) (Figure 1): 1) Parents and offspring were kept at a CO_2_ level of 414 ± 46 μatm (Control treatment); 2) Parents and offspring were kept at control condition, and offspring were exposed to elevated CO_2_ (754 ± 92 μatm) for five days before dissection (Acute treatment); 3) Parents were kept at control condition and offspring were exposed to elevated CO_2_ immediately after hatching for their entire developmental period (Developmental treatment); and 4) both parents as well as offspring were exposed to elevated CO_2_ for the entire duration of the study (Inter-generational treatment). After five weeks juvenile fish were tested for their reaction to CAC as described in Welch and Munday, 2017 with a sample number of 15 individual offspring for each parental behavioural phenotype (T and S) within each of the four CO_2_ treatment (N=120 fish sampled in total). Differences in the percentage of time juveniles spent in CAC was tested by pair-wise comparisons using Tukey’s post hoc tests in R. Six offspring per treatment and phenotype were randomly selected (without prior knowledge of the outcomes of behavioural tests) and dissected for protein expression analysis. Three individuals from each of the tolerant and sensitive group coming from two family lines (Figure 1) were euthanized and the brain was dissected out and snapped frozen in liquid nitrogen and then stored in −80oC for further processing.

**Figure 1.**
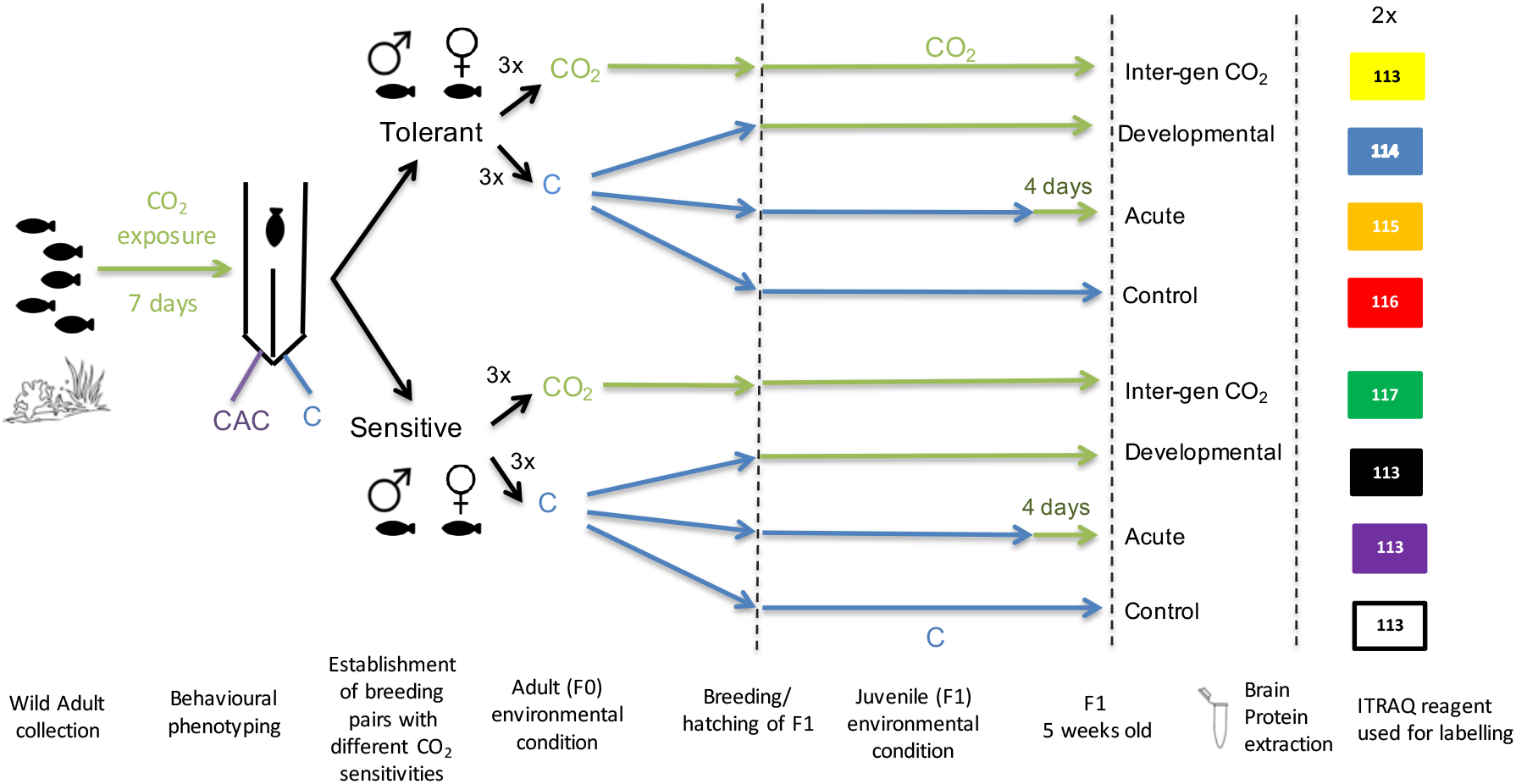
Schematic of experimental design from wild adult fish collection, behavioural testing, environmental CO_2_ exposure treatments and Proteome iTraq experimental design.

### Protein extraction

Proteins from whole brain tissue were extracted along with DNA and RNA with a Qiagen AllPrep DNA/RNA Mini Kit. The flow through at the RNA purification in the manufacturer’s instruction was kept and placed on ice for protein extraction. 3.5 µl of Halt protease & phosphatase inhibitor cocktail 100X, Thermo Fisher Scientific, was added, vortexed and split equally into two tubes and placed on ice for 15 minutes. Then 1000 µl of ice-cold acetone were added and placed on ice for 30 minutes for protein precipitation. Samples were then spun at full speed in a centrifuge at 4oC for 10 minutes. The supernatant of acetone was discarded then without disturbing the protein pellet and the samples were left to dry for 15 minutes and then stored at −80oC.

### Protein digestion and iTRAQ labeling

Dried protein pellets extracted from fish brains were resuspended in 8 M urea lysis buffer with the help of a stick sonicator (Fisher Scientific). Protein concentrations were measured using a 2-D Quant kit (GE Healthcare, UK). Three biological replicates of the same treatment and parental phenotype were pooled at equal concentrations to a final volume of 100 μg. The pools of protein samples were then diluted 1:7 with 50 mM triethylammonium bicarbonate and digested using trypsin (Promega, USA) at an enzyme:protein ratio of 1:40 and left overnight at 37°C. Triflouroacetic acid at 2% was used to inactivate the digestion and peptides were desalted using *Sep*-*Pak C18 cartridges* (100 mg, Water Corporation, USA). Each sample (of three pooled biological replicates) was then labelled with iTRAQ Reagents-8plex reagents (Applied Biosystems) and incubated for 60 minutes. All eight samples (four CO_2_ treatments for two parental phenotypes) were then mixed together and dried by speedvac (Chandramouli et al., 2015). As two family lines were used, two complete iTraq experiments were performed with eight treatment samples (of three pooled biological samples) for a total of 16 treatment samples.

### Peptide fractionation and mass spectrometry analysis

The labeled pool of peptides was fractionated by strong cation exchange chromatography (SCX)^26^ using an Accela 1250 LC system (Thermo Scientific, USA). We obtained a total of 15 peptide fractions and then desalted, dried and resuspended them in 20 μL of LC-MS sample buffer (97% H20, 3% ACN, 0.1% formic acid). The samples were analyzed through three technical replicates using a Q Exactive HF mass spectrometer (Thermo Scientific, Germany) coupled with an UltiMate™ 3000 UHPLC (Thermo Scientific). Introduction of the sample into mass spectrometer was done through a Nanospray Flex (Thermo Scientific) with an electrospray potential of 1.5 kV and the ion transfer tube temperature was set at 160°C. The Q Exactive was set to perform data acquisition in the positive ion mode. A full MS scan (350-1400 m/z range) was acquired in the Orbitrap at a resolution of 60,000 (at 200 m/z) in a profile mode, a maximum ion accumulation time of 100 milliseconds and a target value of 3 × e^6^. Charge state screening for precursor ion was activated. The ten most intense ions above a 2e4 threshold and carrying multiple charges were selected for fragmentation using higher energy collision dissociation (HCD) and the resolution was set as 15000. Dynamic exclusion for HCD fragmentation was 20 seconds. Other setting for fragment ions included a maximum ion accumulation time of 100 milliseconds, a target value of 1 × e^5^, a normalized collision energy at 28%, and isolation width of 1.8.

### Protein identification and quantification

Mascot generic format (*mgf*) files was used to convert the raw MS data in the Proteome Discoverer 1.4 software (Thermo Scientific) and submitted to MASCOT v2.4 (Matrix Sciences Ltd, United Kingdom) for database search against an *A. polyacanthus* brain protein dataset developed in-house from the genome assembly (Schunter et al., 2016). The mass tolerance was set to 10 ppm for precursors, and 0.5 Da for the MS/MS fragment ion with a maximum allowance of one missed cleavage. Variable modifications included 8-plex iTRAQ at tyrosine and oxidation at methionine. The fixed modifications were set to methylethanethiosulfonate at cysteine and lysine, and 8-plex iTRAQ at N-terminal. The MASCOT result files were processed using Scaffold v4.1.1 (Proteome Software Inc. USA) software for validation of peptide and protein identifications with a threshold of 95% using the Prophet algorithm with Scaffold delta-mass correction. iTRAQ label-based quantitation of the identified proteins was performed using the Scaffold Q+ algorithm. The intensities of all labeled peptides were normalized across all runs (Zhang et al., 2010) and individual quantitative data acquired in each run were normalized using the i-Tracker algorithm (Shadforth et al., 2005). Peptide intensity was normalized within the assigned protein. The reference channel was normalized to produce a 1:1 fold change, and the iTRAQ ratios were then transformed to a log scale. P-values were calculated using a paired t-test with significance level of 0.05. We allowed for missing data in one of the three technical replicates and accepted differential expression at a fold-change level of 1.5 (consistent over technical replicates).

### Differential protein expression

We performed two different iTraq experiments including two family lines for differential protein expression among treatments. The final number of proteins varied between 2,100 to 2,300 with a total of 1,862 proteins overlapping among the two iTraq experiments/family lines. Protein quantification was only considered valid if minimum two technical measurements were valid. As we are measuring relative protein expression between samples, we used average values of technical replicates and normalized these against the control replicate for each biological replicate and protein. Each labeled iTraq replicate was compared to the seven other iTraq replicates (each including 3 biological pooled samples) for all commonly measured proteins. Significant differential expression was accepted at a fold change level of 1.5 between two replicates. Differential expression for each treatment comparison was consolidated among both iTraq experiments and was called when both family lines (represented by the two iTraq experiments) showed differential expression for the particular treatment comparison. In cases where only one of the two families showed differentially expression but the other technical replicates in the family have a trend of up/ down regulation, the overall average of all technical replicates within the two families was considered. Hence, the sum of relative protein expression values of all technical replicates was divided by the number of valid technical replicates. The proteins were considered to be differentially expressed when the average of the technical replicates showed 1.5-fold change in expression difference. Functional annotation of detected proteins was performed on the whole list of proteins obtained by using the in-house *A. polyacanthus* protein database and annotated with gene ontology terms in Blast2GO using an FDR significance cutoff of 0.05 (Conesa et al., 2005).

## Results

### Behavioural responses

The behavioural response of 5-week-old *A. polyacanthus* to CAC was variable among individuals within the same CO_2_ treatment (Figure 2). Nevertheless, there were significant differences among CO*2* treatments and parental phenotypes. At control conditions, the average percent of time spent by offspring of tolerant and sensitive parent in CAC was 14.9 and 20.3% respectively. In the acute elevated CO_2_ treatment, offspring of sensitive parents (W= 647, P= <0.001), but not tolerant parents (W= 870.5, P= 0.243) spent significantly more time in CAC compared with controls (Figure 2). For both parental phenotypes, the developmentally and the inter-generationally elevated CO_2_ treated offspring showed a significant difference in behaviour when compared with the offspring in control condition (T Dev: W=155 & P=<0.001; T Inter: W= 334 & P=<0.001; S Dev: W=833.5 & P=<0.001; S Inter: W= 673 & P=<0.001). When comparing directly between parental phenotypes, only the acute elevated CO_2_ treatment resulted in a significant difference with more time spent in CAC for the offspring of sensitive parents (W=563.5, P<0.001).

**Figure 2.**
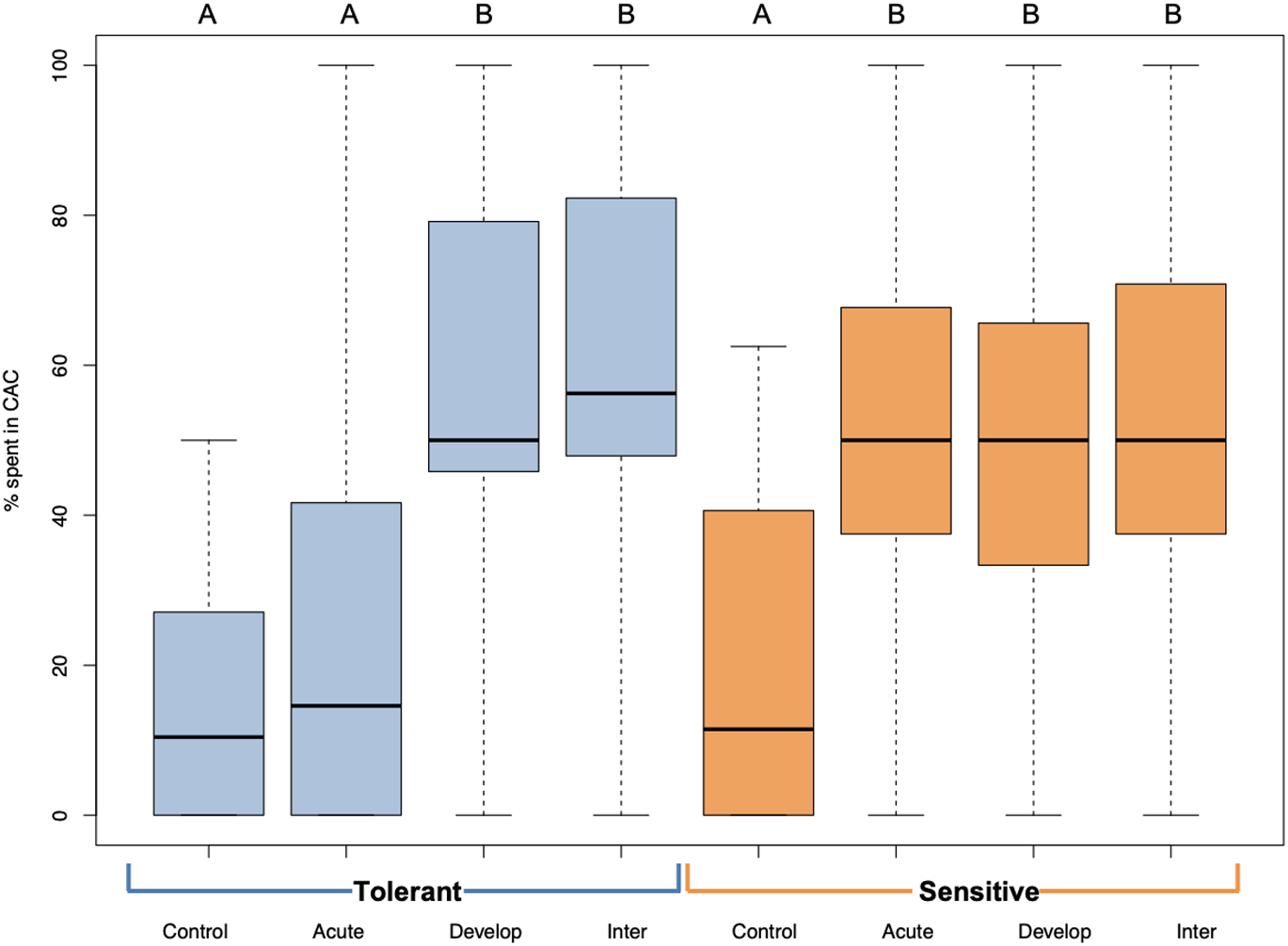
Behavioural responses to chemical alarm cues (CAC) of 5 week old juvenile *A. polyacanthus* exposed to different CO_2_ treatments. Offspring of tolerant parental phenotype are in blue and sensitive in orange. Control, Acute, Develop and Inter are the four different environmental CO_2_ exposures. Different letters represent significant differences between treatments (Tukey, *P* < 0.05). N= 15 for each of the eight treatments - phenotypes combinations.

### Proteome responses to acute elevated CO_2_ conditions

Over 1,800 proteins were detected and commonly expressed among the two-family lines. We first considered the offspring of the two parental phenotypes separately to understand the reaction to the different CO_2_ treatments against the respective control individuals. We detected between 0 and 40 differentially expressed proteins among the pair-wise comparisons of different elevated CO_2_ exposures (Figure 3). Individuals exposed to acute elevated CO_2_ for five days exhibited a total of 17 and 40 differentially expressed brain proteins in comparison to the control group for the sensitive and tolerant parental group respectively (Figure 3). Fifteen proteins were found commonly expressed in both groups, representing 35% and 82% in tolerant and sensitive groups respectively, showing a common response to acute exposure to elevated CO_2_ (Table 1). All of the 15 proteins were expressed at higher levels in the acute treatment than in the control condition. One of these proteins was B-cell receptor-associated protein 31-like isoform 1 (BCAP31), which is involved in the expression of MHC class I protein and the negative regulation of GABAergic synapses density (Elmer and McAllister, 2012). Three up-regulated proteins are involved in histone modifications. This includes the non-histone chromosomal protein HMG-14A-like isoform X2 (HMGN2) which is involved in phosphorylation inhibition of nucleosomal histones H3 and H2A; chromobox protein homolog 1-like (CBX1) which is associated with histone binding; and prothymosin alpha-B-like (PTMAB) which plays a role in histone methylation.

**Table 1.**
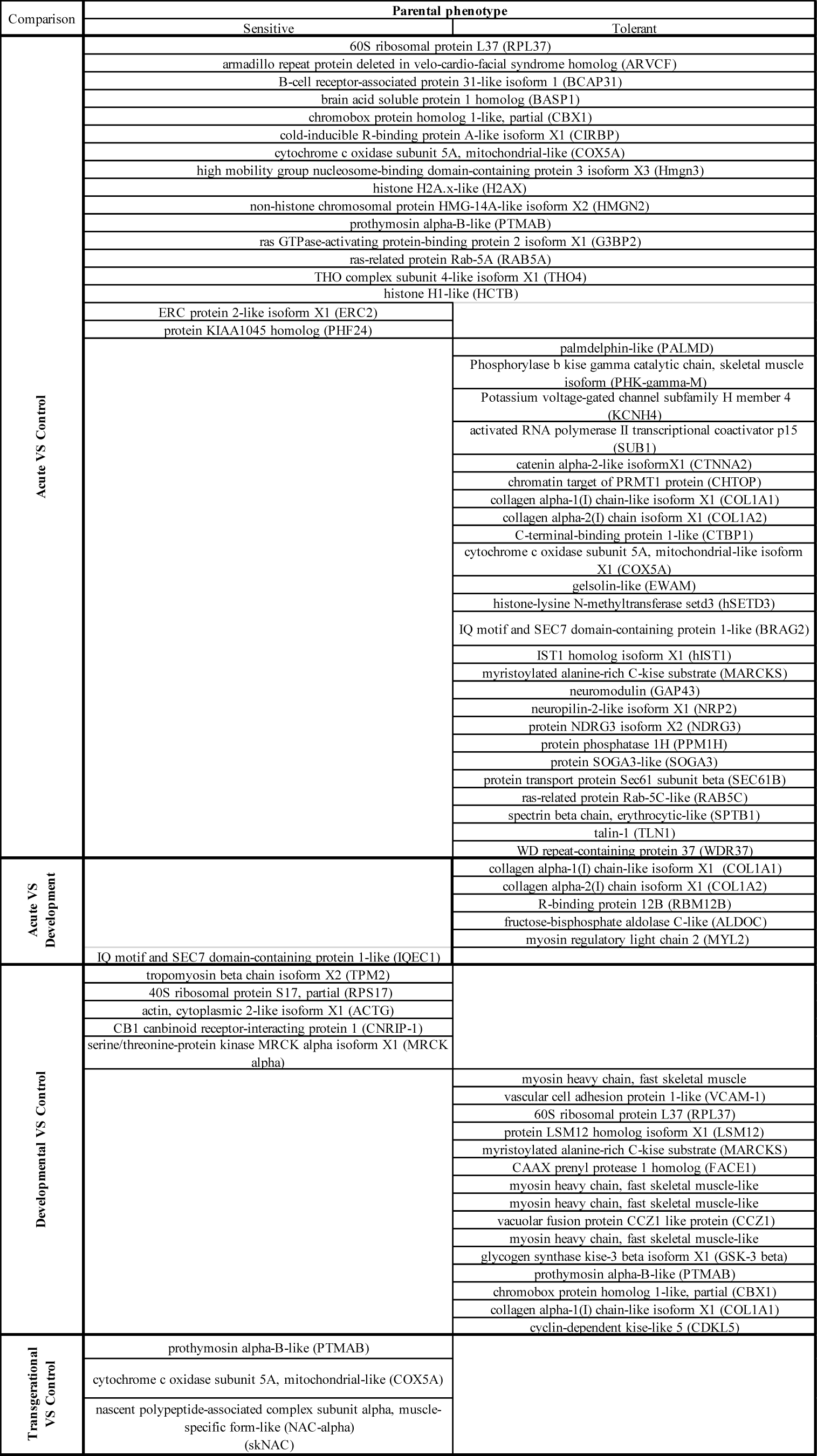
Differentially expressed proteins (DEPs) in the brain of 5 week old juvenile *A. polyacanthus* exposed to four different CO_2_ treatments: Acute vs Control, Developmental vs Control and Inter-generational vs Control. Differential expression was performed separately for each parental phenotype against the respective control individuals. Top rows in the table show DEPs for each comparison significantly differentially expressed for offspring of both parental phenotypes.

**Figure 3.**
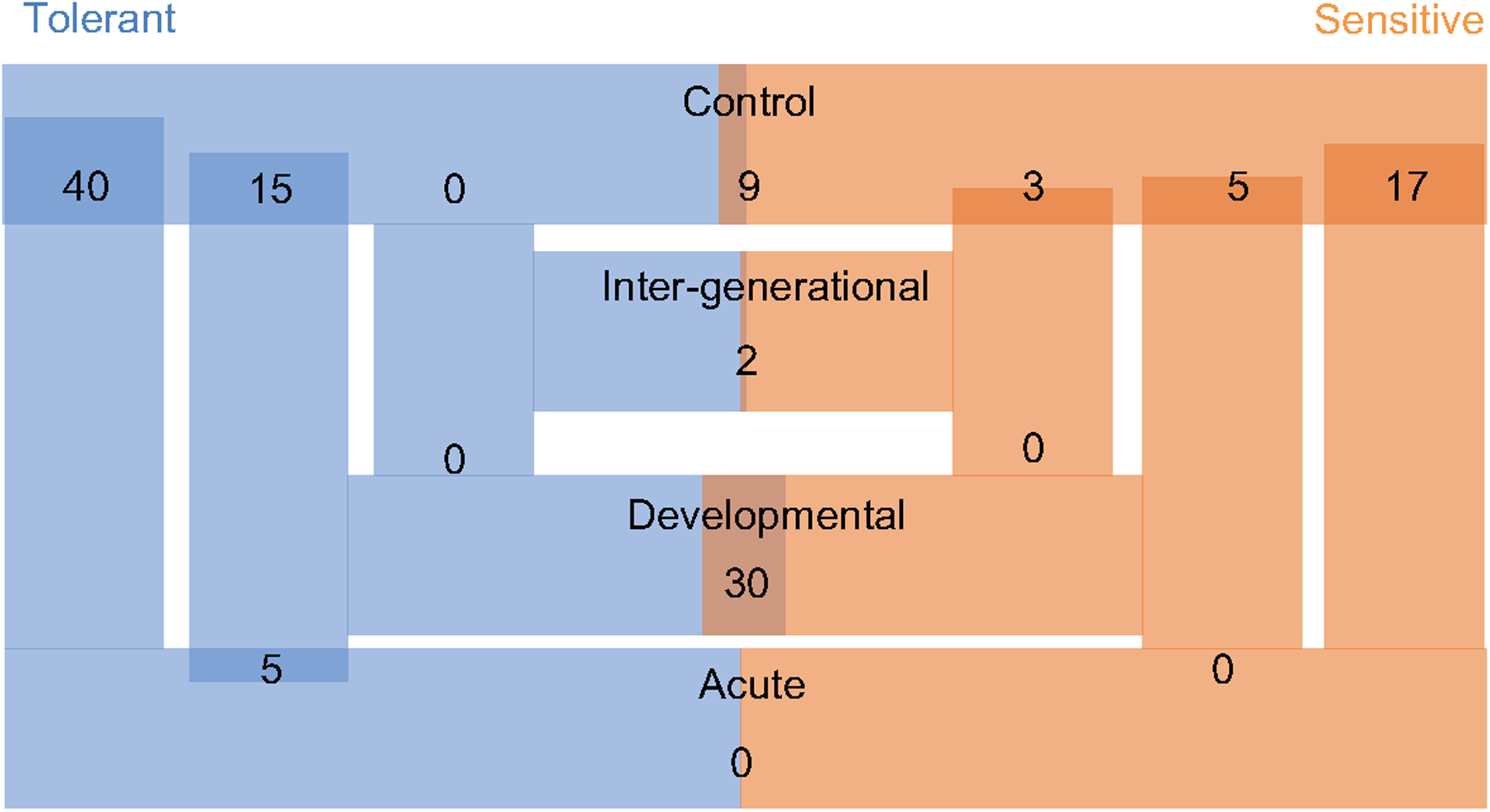
The experimental conditions and the number of differentially expressed proteins in the. ‘Tolerant’ and ‘Sensitive’ refers to the behavioural parental phenotypes. Control, Acute, Developmental and Inter-generational refer to each of the experimental CO_2_ conditions (see Figure 1).

### Proteome responses to longer-term CO_2_ exposure

The longer-term developmental elevated CO_2_ exposure resulted in a less pronounced protein response when compared with the respective control individuals, with fewer differentially expressed proteins than in the acute CO_2_ treatment (Figure 3). There was also no overlap in the differential expression among the offspring of sensitive and tolerant parents for this CO_2_ treatment (Table 1). In the offspring of sensitive parents, five proteins were differentially expressed with one expressed at a higher level and four at a lower level, including CB1 cannabinoid receptor-interacting protein 1 (CNRIP-1), which is related to synaptic plasticity. Fifteen proteins were differentially expressed in the offspring of tolerant parents, none of which were differentially expressed in the offspring of sensitive parents (Figure 3 & Supplementary Table S1). Two upregulated proteins, vacuolar fusion CCZ1 like protein (CCZ1) and vascular cell adhesion protein (VCAM-1), are functionally related to calcium ion binding and organ development respectively (Supplementary Table S1). 60S ribosomal protein L37 (RPL37), chromobox protein homolog 1-like (CBX1) and prothymosin alpha-B-like (PTMAB) in the offspring of tolerant parents showed similar down-regulation patterns as seen for fish in the acute CO_2_ treatment. Further proteins found to be differentially expressed between the more longer-term developmental treatment and the acute exposure are several collagen alpha chain isoforms (COL1A1 & COL1A2; Table as well as myosin regulatory chain 2 (MYL2) for the offspring of sensitive parents whereas only one protein was differentially expressed for the offspring of tolerant parents (IQ motif and SEC7 domain-containing protein).

The acute and developmental CO_2_ treatments reflected the within-generational responses. In the offspring of sensitive parents, 17 proteins were differentially expressed in the acute CO_2_ condition while only five were differentially expressed in the developmental CO_2_ condition and none overlap between the two comparisons (Figure 3, Supplementary Table S1). As for the offspring of tolerant parents, five proteins were commonly differentially expressed for the acute and developmental CO_2_ condition against the control condition which include functions such as immune response and epigenetic control (RPL37, PTMAB, MARCKS, CBX1 and COL1A1; see Supplementary Table S1).

### Inter-generational exposure to elevated CO_2_

To determine the effects of parental CO_2_ exposure on the offspring brain protein expression, we compared the inter-generational CO_2_ treatment with the control condition for both parental groups. No proteins were found to be differentially expressed in the tolerant group, whereas three proteins were differentially expressed in the sensitive group (Table 1). The two down-regulated proteins are prothymosin alpha-B-like (PTMAB) and cytochrome c oxidase subunit 5A mitochondrial-like protein (COX5A). The third (up-regulated) protein, nascent polypeptide-associated complex subunit complex alpha muscle specific form-like protein (NAC-alpha), is functionally related to the TBP-class protein binding.

### Differential responses due to parental phenotype

In our experiment we were also able to directly compare differential brain protein expression between the offspring of the two parental phenotypes in the same environmental treatment (Figure 3). In control condition, nine proteins differed significantly in expression between offspring of sensitive and of tolerant parents of which five were expressed at lower levels and four at higher levels for the offspring of sensitive parents in comparison to the offspring of tolerant parents (Supplementary Materials Table S2). Both parental phenotypes reacted similarly in the acute elevated CO_2_ treatment as there was no significant difference in protein expression directly between the offspring of the two parental phenotypes for this environmental condition (Figure 3). The similar response between offspring of tolerant and sensitive parents in the acute elevated CO_2_ treatment is also shown by the previously described overlap of differentially expressed proteins when comparing between environmental conditions of acute and control CO_2_ (Table 1). By contrast, 30 proteins were differentially expressed between the offspring of the two parental phenotypes in the developmental elevated CO_2_ treatment. The majority of the differentially expressed proteins (87%) were expressed at a higher level in the offspring of tolerant parents (Supplementary Table S2). As for the inter-generational CO_2_ condition, only two proteins were differentially expressed between the offspring of the two parental phenotypes. The function of one of the proteins is not known. The other protein was AMP deamise 2-like isoform X1 protein (AMPD2), which is functionally related to the cyclic purine nucleotide metabolic process and energy homeostasis.

The different responses on the brain protein level for offspring of both parental phenotypes as well as for the different environmental treatments can be visualized in four recurring differentially expressed proteins, including ribosomal protein L37 (RPL37), chromobox protein homolog 1-like (CBX1), cytochrome c oxidase subunit 5A (COX5A) and prothymosin alpha-B-like (PTMAB; Figure 4). In the acute elevated CO_2_ treatment, all four proteins were down-regulated in offspring of both sensitive and tolerant parental phenotype (Figure 4a-d). However, in the developmental elevated CO_2_ treatment, the proteomic responses of the offspring from the two parental phenotypes diverged. RPL37 and CBX1 were down-regulated in the offspring of tolerant parents, whereas the expression of these proteins was at control levels in offspring of sensitive parents (Figure 4a-b). The diverged proteomic responses between the two parental phenotype groups were further detected in the inter-generational elevated CO_2_ treatment. COX5A and PTMAB were down-regulated in the offspring of sensitive parents (Figure 4c-d), whereas the expression of these proteins was at control levels in the offspring of tolerant parents.

**Figure 4.**
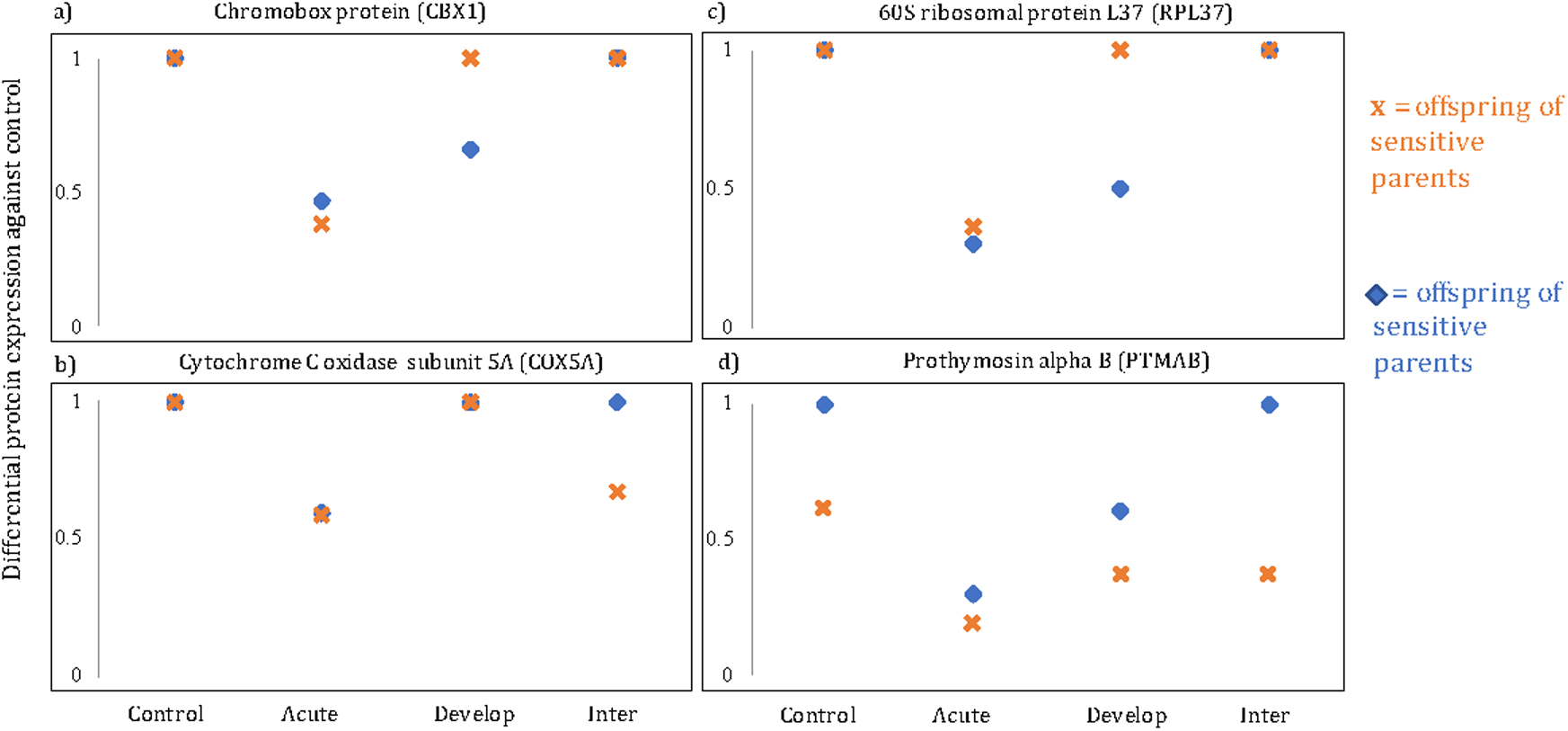
Level of protein expression in the sensitive and tolerant group, a) chromobox protein homolog 1-like (CBX1), b) cytochrome C oxidase subunit 5A, mitochondrial-like (COX5A), c) 60S ribosomal protein L37 (RPL37) and d) prothymosin alpha B-like (PTMAB). Control, Acute, Develop, Inter refer to the control, acute, developmental and inter-generational treatment respectively.

## Discussion

Measuring brain proteome expression of organisms exposed to elevated CO_2_ conditions for different durations, within and between generations, can provide insight into how ocean acidification affects biological processes over relevant timescales. In this study we tested the effects of elevated CO_2_ treatment and parental phenotype on proteins expression in the brain of five-week old developing juvenile *Acanthochromis polyacanthus*. Protein expression varied greatly among the different CO_2_ treatments, as well as between offspring from the two parental phenotypes, revealing treatment-specific responses to ocean acidification conditions in the brain proteome. This is consistent with previous results on the transcriptional level, where responses largely depended on the fish’s life stage and the duration of the exposure (Schunter et al., 2018).

The acute (5 day) elevated CO_2_ treatment resulted in the largest change in brain protein abundance of all treatments. Acutely treated fish were also the only group with a partial overlap in proteins response between the parental phenotypes. This reveals a large and partially common stress response to an acute change in environment CO_2_. We found changes in the expression of stress-related proteins, for example the cytochrome c oxidase subunit 5A (COX5A) which regulates cell activation and differentiation (Tarasenko et al., 2017). Studies in fish have shown that highly oxidative organs such as the brain depend on cytochrome c oxidase for normal functioning as it serves as a marker for neuronal activity (Wong-Riley, 2012) and Cox deficient fish diplayed neurological abnormalities (Diaz, 2010). Another stress-related response in the acute treatment was the differential regulation of proteins involved in epigenetic regulation. Amelioration of environmental CO_2_ changes by epigenetic regulation might be seen in our juvenile fish through the downregulation of several proteins linked to histone regulation. We found elevated proteins expression in histone H1-like, together with PTMAB which binds to linker histone H1 (Karetsou et al., 1998; Zakharova et al., 2011), which also exhibited higher transcriptional levels in brains of five-month-old *A. polyacanthus* with elevated CO_2_ exposure (Schunter et al., 2016). The differential regulation of the genome, including silencing, enhancing and splicing of genes, relies heavily on these histone modifiers (Bird, 2002) and has been shown to enable plastic responses to environmentally induced changes (Turner, 2009). The regulation patterns of epigenetic markers at different life stage reveals an important molecular mechanism by which individuals respond to acute environmental change in CO_2_ conditions.

In contrast, during the longer-term developmental exposure to elevated CO_2_ the functions of differentially expressed proteins were related to neurological development in juvenile *A. polyacanthus*. The CB1 cannabinoid receptor-interacting protein 1 (CNRIP1), for example, is a protein involved in synaptic plasticity and modulates neurotransmission (Howlett et al., 2010) and was differentially expressed in the offspring of parents that were behaviourally sensitive to elevated CO_2_. Previous studies on zebrafish revealed that CNRIP1 is highly expressed in the nervous system, particularly in developing fish, and CNRIP1 interacts with the cannabinoid receptor 1, a receptor critical for the functional development of vision and locomotion (Fin et al., 2017; Oltrabella et al., 2017). In the offspring of tolerant parents, we further found differentially expressed proteins, such as the vascular cell adhesion protein (VCAM-1), to be involved in cellular process and organ development (Kong et al., 2018). Between the acute and developmental treatments we also detected differentially expressed proteins that are involved in connective tissue, such as collagen, which is important in brain development as well as brain repair and nerve regeneration (Gregorio et al., 2018). The myosin regulatory light chain 2, also differentially expressed in the brain of fish in the developmental elevated CO_2_ treatment (in comparison to acute and control) is involved in neuronal development and this protein was also found differentially expressed after exposure to a neurotoxin in medaka fish (Tian et al., 2011). Our results suggest that the neurological pathways and structure in the brain could be affected in particular when exposed to elevated CO_2_ during development.

Few proteins were differentially expressed when offspring experienced developmental exposure to elevated CO_2_ in combination with previous exposure of the parents to elevated CO_2_, revealing an effect of parental exposure on the proteome of the offspring. Only two proteins, prothymosin alpha-B-like (PTMAB) and cytochrome c oxidase subunit 5A (COX5A), were up-regulated independent of parental exposure, but only in the offspring of sensitive parents. Similarly, in the European seabass an immune gene stayed upregulated in the olfactory epithelium despite previous parental exposure to ocean acidification (Mazurais et al., 2020). However, in our study all other differentially expressed proteins in the within-generational treatments (acute and developmental) were at control levels in the inter-generational treatment. This hints to a non-genetic effect exerted by parents on the offspring through maternal provisioning and/or epigenetic inheritance (Bonduriansky et al., 2012; Salinas et al., 2013). Such parental effects have widely been recognized and has, in some cases, been directly proven to increase offspring’s fitness (Evans et al., 2014). The molecular adjustments in fish exposed to ocean acidification conditions are of lesser magnitude when their parents were also exposed to elevated CO_2_, both on the transcriptional level (Schunter et al., 2018) as well as seen here on the protein level. Despite this influence of previous parental exposure to elevated CO_2_ on the proteome, behavioural responses to chemical alarm cues (CAC) remained impaired in the inter-generational treatment. This is consistent with a previous study on the effect of elevated CO_2_ on behaviour, based on a larger sample size, where previous parental treatment revealed no difference in the reaction to CAC (Welch and Munday, 2017). Hence, we see a change in the proteome response of juvenile damselfish with parental exposure, but this is not reflected in the behavioural response. Inter-generational acclimation might therefore happen on the molecular level in the brains of *A.polycanthus*, but this does not alleviate the behavioural impairments to ocean acidification.

Previous studies have linked behavioural impairments in fish under elevated CO_2_ conditions to the altered functions of GABA neurotransmitter receptor (Nilsson et al., 2012; Schunter et al., 2019). One previous study in *A. polyacanthus* found highly up-regulated expression of GABA related genes in acute and developmental exposure to elevated CO_2_ (Schunter et al., 2018). In our study of five-week-old fish, there were no differentially expressed proteins related to GABA receptor functions. Our experiment recovered a subset of approximately 2,000 proteins, in which we did detect gamma-aminobutyric acid receptor-associated proteins and aldehyde dehydrogese family 9 member A1-B-like protein, previously found to be differentially expressed on the transcript level in older fish (Schunter et al., 2016); however in our 5-week old fish brains these proteins were not differentially expressed. Our behavioural trials showed that after acute CO_2_ exposure our five-week old fish differed in their behavioural responses to CAC between the two different parental phenotypes. Despite this difference in behaviour, these fish exhibited no differentially expressed proteins, when directly compared, revealing that under short-term elevated CO_2_ fish of both parental phenotypes exhibit some similarity in their reaction at the brain protein level. For any other CO_2_ treatment (developmental or inter-generational), we do find distinct protein expression patterns between offspring from the different parental phenotypes. Hence in our five-week old fish the signal found in the brain proteome and the behaviour does not show a highly concordant pattern. The connection between impaired behaviour and protein responses to environmental elevated CO_2_ therefore remains unclear and it is possible that the brain proteins expression is not representative of behavioural patterns due to the extended time needed for protein synthesis. Nevertheless, our results show that: 1) developing juveniles are affected in their brain proteins expression by near-future elevated CO_2_ conditions, 2) that the impact on protein expression depends on the length of exposure to elevated CO_2_ exposure, with fewer differentially expressed proteins being exhibited in longer-terms and inter-generational treatments, and 3) that variation of parental sensitivity to behavioural effects of elevated CO_2_ affect the brain proteome of juveniles, especially under developmental exposure to elevated CO_2_. This variation in molecular response in a juvenile fish reveals a wide repertoire of plastic and inter-generational responses to elevated CO_2_ which could aid a coral reef fish to adjust to near-future ocean acidification conditions.

## Acknowledgments

This study was supported by the Australian Research Council (ARC) and the ARC Centre of Excellence for Coral Reef Studies (P.L.M), the Office of Competitive Research Funds OCRF-2014-CRG3-62140408 from the King Abdullah University of Science and Technology (T.R., P.L.M., C.S.). This project was completed under James Cook University (JCU) ethics permit A1828. We thank the Marine and Aquaculture Research Facilities Unit (JCU), the Schunter lab members at the Swire Institute of Marine Science (SWIMS) and Biosciences Core Laboratory (KAUST) for support and assistance.

## Author contributions

M.W. and P.L.M designed and managed the fish rearing experiments. M.W. performed the fish behavioural phenotyping. C.S. extracted the proteins, designed the protein expression experiment, prepared the i-TRAQ libraries and identified the proteins. H.H.T & C.S. analyzed protein expression data and interpreted the results. H.H.T., C.S., P.L.M. and T. R. wrote the paper and all authors read and approved the final manuscript.

